# Chromatin state modeling across individuals reveals global patterns of histone modifications

**DOI:** 10.1101/2022.08.02.502571

**Authors:** Jennifer Zou, Jason Ernst

## Abstract

Epigenetic mapping studies across individuals have identified many positions of epigenetic variation in various human tissues and conditions. However the relationships between these positions, and in particular global patterns that recur in many regions of the genome remains understudied. In this study, we use a stacked chromatin state model to systematically learn global patterns of epigenetic variation across individuals and annotate the human genome based on them. We applied this framework to histone modification data across individuals in lymphoblastoid cell lines and across autism spectrum disorder cases and controls in prefrontal cortex tissue. We find that global patterns are correlated across multiple histone modifications and with gene expression. We used the global patterns as a framework to predict transregulators, identify trans-QTL, and study complex disease. The frameworks for identifying and analyzing global patterns of epigenetic variation are general and we expect will be useful in other systems.

## Introduction

Understanding molecular variation is fundamental to understanding variation in complex traits. Many studies have identified molecular variation across individuals in transcription factor (TF) binding, gene expression, histone modifications, and other molecular phenotypes [1, 2, 3, 4, 5, 6, 7].

Understanding variation in histone modifications that are associated with enhancers or promoters can be of particular interest since variants for many diseases and phenotypes are enriched in enhancer and promoter regions of the genome [8, 9, 10, 11, 12]. A number of studies have mapped histone modifications associated with enhancers and promoters across many individuals [13, 3, 4, 5, 14, 15, 16, 6]. These studies have identified thousands of regions where histone modifications differ across individuals.

These previous studies often identify a set of consensus regions across individuals with histone modification signal, such as merged peaks, and perform a marginal association test between each region to an external data set. For example, in histone quantitative trait loci (hQTL) studies, variation is identified by associating histone modification signal across individuals with genetic variants. Similarly, for differential peak analysis, variation in a single region is associated with an external label, such as cases and controls [4, 17, 18, 19]. Another approach, which allows for joint analysis of multiple marks is to learn combinatorial and spatial patterns of epigenetic marks that are associated with distinct biological functions (“chromatin states”) [20]. This approach has been previously applied to histone modification data across multiple individuals by virtually concatenating data across individuals for each data type and learning a chromatin state model using the ChromHMM software [21, 3]. Using the learned model, individual-specific genome chromatin state annotations were generated, which contain chromatin state assignments for each individual genome-wide. These annotations were then used to identify regions with variable chromatin states across individuals [3]. While informative, one understudied aspect of these previous approaches is the relationships between the variable regions, in particular recurring patterns of epigenetic variation across individuals observed in many regions of the genome.

One biological reason we may expect to observe recurring epigenetic patterns across individuals is that a transcription factor may have differential activity across individuals. This could be reflected in corresponding differential activity of histone modifications at many of its binding locations across the genome. Such transcription factors can act as “trans-regulators” potentially affecting the expression of many genes in the genome. Reflecting the importance of trans-regulation, it has been estimated that 60-75% of the heritability in gene expression is explained by distal effects [22, 23, 24, 25]. However, the identification of trans-regulators and genetic variants associated with them through trans-expression QTL (trans-eQTL) mapping is challenging. This is the case compared with cis-expression QTLs (cis-eQTLs) because of the much larger number of association tests that need to be performed, which results in lower statistical power [26]. The recurring patterns of global variation have the potential to provide useful information towards the challenge of identifying trans-regulators and trans-eQTLs in a given cell type.

In this study, we use a “stacked” chromatin state model to systematically learn global patterns of epigenetic variation across individuals and annotate the genome based on them [21, 27, 28]. The “stacked” chromatin state model is based on a multivariate hidden Markov model (HMM) that learns combinatorial and spatial patterns across multiple individuals of one or more marks that recur in many regions of the genome. We first develop and use the global patterns to predict transregulators and identify trans-QTLs in lymphoblastoid cell line. Then, we demonstrate how this framework can be applied to histone modification data from autism spectrum disorder (ASD) cases and controls in prefrontal cortex tissue. While previous studies have identified numerous molecular features, including RNA expression, RNA splicing, and histone modifications, that differ between ASD cases and controls [29, 30, 31, 4, 32, 33], we apply the global patterns framework and show that global patterns are also associated with diagnosis status. We expect identifying global patterns of epigenetic variation will be a useful framework to study transcriptional regulatory networks and complex disease in other systems.

## Results

### Systematic genomic annotation of chromatin variation across individuals

We learned a stacked ChromHMM model where all histone marks in all individuals are used as features by applying a “stacked” version of the ChromHMM framework [21, 27, 28]. We used genome-wide histone modification data quantified in 200bp non-overlapping bins across multiple individuals and marks. We first regressed out the effects of known confounders before training the model (Methods). Similar to the ChromHMM framework when data from only a single cell type is used to train a model, we binarized the data using a Poisson background model and used this as input to ChromHMM [20, 21]. Unlike the standard use of ChromHMM, in this framework, each hidden state learned corresponds to a combinatorial and spatial pattern across marks and individuals, which we call a “global pattern.” The emission probabilities correspond to the probability of observing a mark in a specific individual given a global pattern. After the global patterns are learned, we annotate the genome at 200bp resolution with the most likely hidden state of the HMM.

### Learning global patterns in lymphoblastoid cell line

We first applied the stacked ChromHMM model to a data set of 75 individuals with 3 marks (H3K27ac, H3K4me1, H3K4me3) in the lymphoblastoid cell line (LCL) [15]. Different combinations of these three marks within a single individual are associated with different types of promoters and enhancers [34, 12, 15, 35]. We trained stacked ChromHMM models to learn global patterns across both individuals and marks (Methods, Figure 1). We learned models with between 5 and 100 states in increments of 5. We used the models learned to then segment and annotate the genome according to these global patterns.

**Figure 1:**
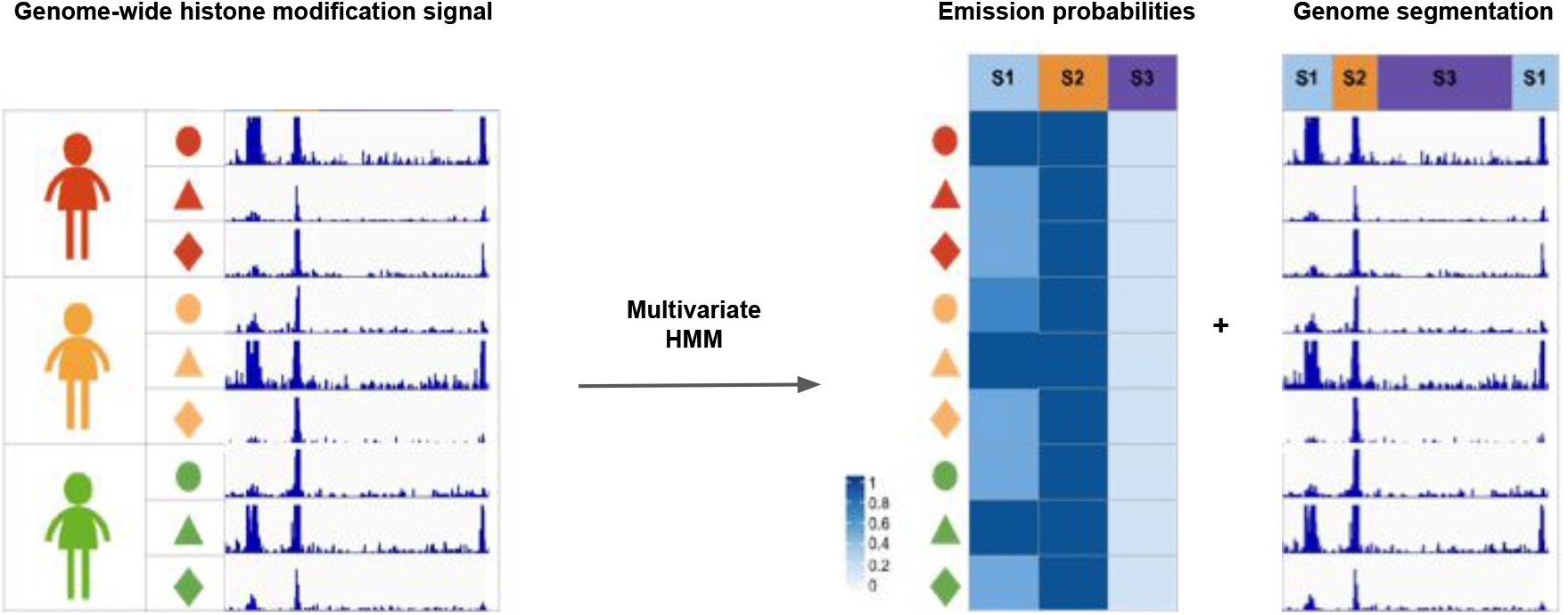
Method overview. We trained a stacked ChromHMM model using genome-wide histone modification signal from multiple individuals and marks (“Genome-wide histone modification signal”). This model learns global patterns of epigenetic variation that recur in many regions of the genome (“Emission probabilities”). The model learning is agnostic to the mark labels. The emission probabilities learned correspond to the probability of observing signal from each data set conditioned on being in each hidden state. We used the model learned to annotate the genome according to these patterns at 200bp resolution (“Genome segmentation”).

As an initial validation of the models we tested whether the global patterns were internally consistent across pairs of histone modifications, even though they are from models that were learned agnostic to mark labels during training. Based on prior knowledge that pairs of these three marks frequently co-occur in the genome, we expect that the emission parameters for pairs of marks to be correlated for many of the states. Specifically, we calculated the median Spearman correlation of emission parameters for pairs of marks across individuals for each global pattern as a function of the number of states in the model. The median pairwise correlation between marks increased rapidly until the number of hidden states was increased to 35. Pairs of histone modifications corresponding to active promoters (H3K4me3 and H3K27ac) and enhancers (H3K4me1 and H3K27ac) remained high (>0.5) for models with up to 100 states (Figure S1). High correlations were observed even though the models were agnostic to the mark and individual labels in the training process. This suggests that a global pattern is less likely to be caused by technical issues with the ChIP-seq experiments and more likely to be associated with differences at the sample level.

### Common genetic variation associated with LCL global patterns

To investigate the genetic basis of global epigenetic patterns across individuals, we performed what we termed global pattern quantitative trait association analysis. For each model that we had learned, we associated common variants with minor allele frequency (MAF) greater than 0.05 in the data set [15] with the emission parameters of each global pattern to identify significantly associated (*p*_*adj*_ < 0.05) global pattern quantitative trait loci (“gQTLs”). The number of gQTLs was maximized with the 85-state model, (2948 gQTLs, Figure S3), which we selected for further analysis. In this 85-state model, 38 states were associated with at least one gQTL. We also verified that global patterns for the 85-state model were robustly learned across different subsets of the genome (Median Spearman correlation of emissions = 0.93, Methods, Figure S2).

We then sought to understand whether the gQTLs identified were biologically relevant for the LCL cell type by performing a GREAT gene set enrichment analyses for the gQTLs [36]. GREAT analyzes genes near a set of genomic regions and tests for ontology and phenotype enrichments compared to a background set of regions. We used the gQTLs obtained as the foreground and the whole genome as the background. The “regulation of leukocyte cell-cell adhesion”, “regulation of lymphocyte activation”, and “regulation of T cell activation” gene ontology (GO) terms were significantly enriched for the gQTLs (FDR < 5%, Supplementary Data 1). These enrichments are expected since the lymphoblastoid cell line is derived from lymphocyte cells, which are critical for immune system functions. While we also observed significant enrichment for terms not directly related to immune function, we do not expect all global patterns to be cell type-specific. Additionally, there may be pleiotropy between immune function and other complex traits. Enrichment of terms relevant to immune function suggest that some gQTLs are biologically relevant for the LCL cell type and that global patterns may be informative for identifying sources of molecular variation associated with immune function.

### LCL global patterns enriched for active regions of the genome

To characterize the types of genomic regions found in the global patterns learned in the 85-state model, we computed overlap enrichments for previous chromatin state annotations in the LCL data obtained (Methods, Figure S4). These chromatin states annotations correspond to combinatorial and spatial patterns of epigenetic datasets within LCLs for a single individual and have been previously given candidate biological descriptions. We identified the chromatin state with the highest fold enrichment among states significantly enriched (overlap enrichment > 1 and Binomial Test, FDR<5%; Methods) for each global pattern. The majority of global patterns are most highly enriched for enhancer and promoter chromatin states, which is expected given the histone modifications used to learn the model (Figure 2A). We define promoter-like global patterns to be global patterns with the highest enrichment for promoter states and enhancer-like global patterns to be those with the highest enrichment for enhancer states from the reference chromatin state annotations.

**Figure 2:**
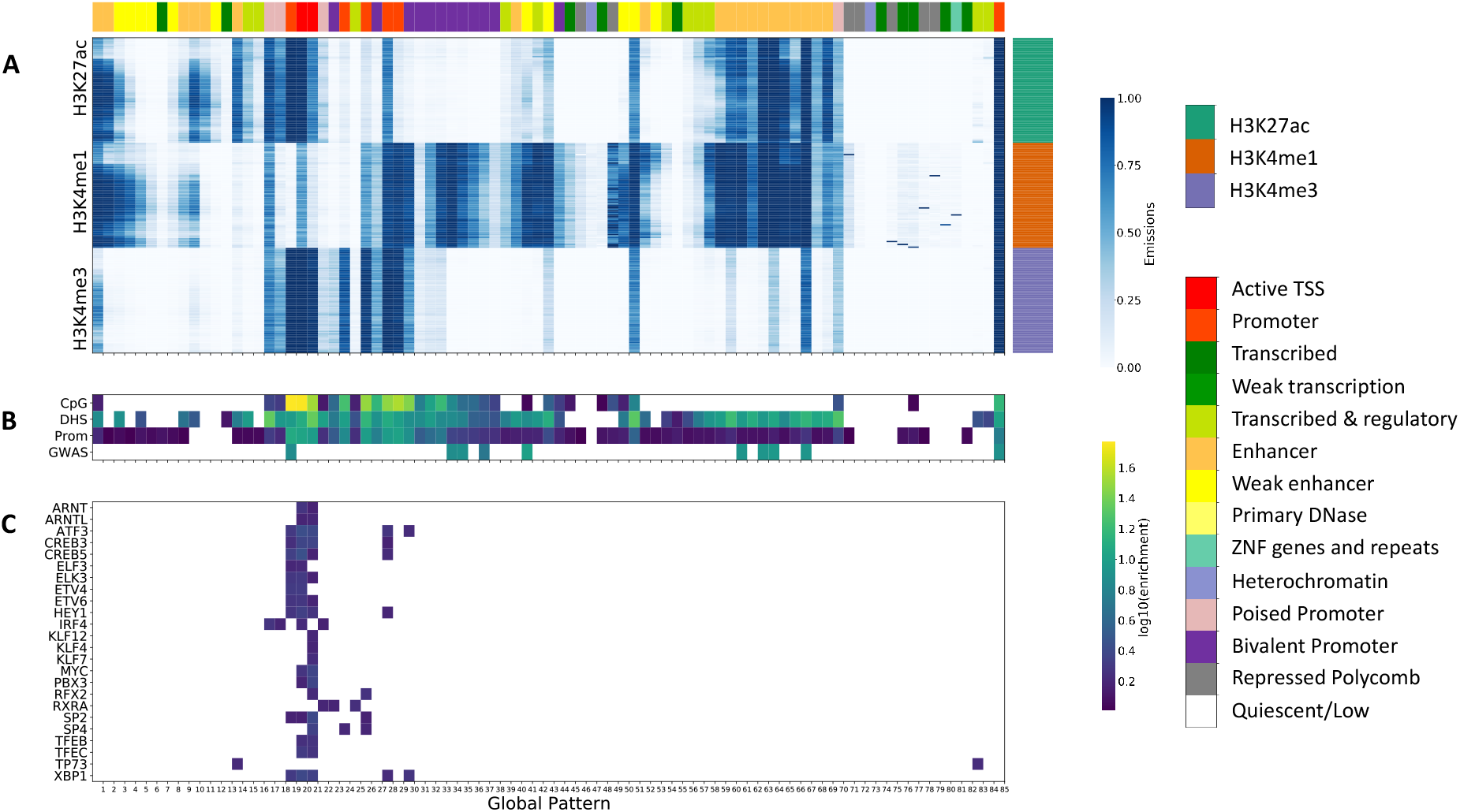
LCL 85-state model. A) Emission matrix for the 85-state LCL model. The x-axis (“Global Pattern”) shows the global patterns learned, and the y-axis shows the datasets, which are grouped by the indicated histone modifications. The ordering of individuals is the same for each histone modification. Each global pattern is annotated (top) with a previous LCL chromatin state annotation for one individual [40] with the highest significant enrichment. B) Overlap enrichments for CpG Islands [37], consensus DNase I hypersensitive sites [15] measured in the same samples, and promoter annotations computed from GENCODE transcription start site (TSS) annotations [38], and immune-related GWAS variants [9]. Only significant enrichments are shown (Binomial Test, FDR<5%, fold > 1). C. Motif enrichments for 24 TFs with motifs enriched in at least one global pattern (FDR<5%, log10(fold enrichment) > 1.5) and gene expression associated with global patterns (FDR<5%). Only significant enrichments are shown (Fisher’s Exact Test, FDR<5%, log10(enrichment) > 1.5).

We also evaluated global pattern enrichments for other external annotations that were not based on histone modifications. These included CpG Islands [37], DNase I hypersensitive sites [15], and promoter annotations defined as regions 2KB from GENCODE transcription start sites (TSS) [38], which as expected showed significant enrichments for many global patterns (Figure 2B, Binomial Test, FDR<5%). CpG Islands tend to be enriched in promoter-like global patterns, whereas DNase I hypersensitive sites are enriched in both promoter-like and enhancer-like global patterns. We also evaluated enrichments for fine-mapped GWAS variants for 39 immune-related diseases [9], which revealed nine global patterns of histone modifications with significant enrichments (Figure 2B, Binomial Test, FDR<5%). These GWAS enriched global patterns were strongly enriched for promoters and enhancers, which is consistent with previous analyses [9, 12, 39].

### LCL global patterns associated with large-scale molecular variation

In addition to using external annotations of the human genome, we associated the global patterns with external gene expression [15] and protein quantification [41] data sets across individuals, which were not used to train the models.

We associated the global patterns with the expression of genes in a subset of the individuals (*n* = 54) for which gene expression data was available. We identified 4774 genes with expression patterns significantly associated with at least one global patterns (Linear model, FDR<5%), with 38 global patterns represented among these associations, indicating that the global patterns co-vary with the expression of a large number of genes. We also computed the average correlation between each global pattern and gene expression at different distances to the global pattern in the annotation (for each 100bp window, up to 500KB upstream and downstream). We found that global patterns tend to more highly associate with genes in relatively closer proximity (i.e, 100KB, Figures S5-S7), as expected.

We also associated the global patterns with protein abundance data for a subset of 60 out of the 75 individuals for which the protein data was available [41]. Of the 60 individuals with both protein abundance data and histone modification data, 44 also had expression data. After intersection of the protein data with their corresponding gene expression data [15], there were 4371 genes with protein abundance data and gene expression data. We correlated the abundance of each protein across individuals with the emission parameters for each global pattern (Methods). We identified 594 proteins that were significantly associated with at least one global pattern (Linear model, FDR < 5%). Of these 594 proteins, we identified 258 proteins that also had differential expression significantly associated with at least one global pattern. Of these 258 proteins, 143 were associated with at least one of the same global patterns as its corresponding gene expression values. We note the partial agreement with corresponding differential gene expression could be explained by a number of factors. First, of the 4774 genes with expression patterns significantly associated with a global pattern, 3425 did not have protein data available. Second, the number of samples differed between the data sets with only 44 samples shared between protein abundance, histone modifications, and gene expression. Since global patterns reflect different subsets of individuals with histone modification signal, lack of overlap could account for the partial agreement. Finally, proteins with differential expression but not differential protein levels could potentially be explained by post-transcriptional differences between individuals [41, 42].

Although not all genes with expression levels associated with global patterns also have protein levels associated with global patterns, there is greater overlap between these two sets of genes than expected by chance (Permutation test, p<1e-4). These results indicate that LCL global patterns based on chromatin are associated with large-scale molecular variation across individuals as determined by other assays further supporting their biological relevance.

### Predicting trans-regulators in LCL

As the global patterns of histone modifications might be associated with TFs that have differential activity across individuals, the identification of enriched DNA motifs at genomic positions annotated to specific patterns can suggest trans-regulators associated with them. Thus, in order to identify potential transregulators, we first performed TF motif enrichment analysis for each global pattern. We calculated motif enrichment of 602 TF motifs in the ENCODE motifs database [43] compared to shuffled motifs. We observed enrichment of motifs for 20 of the 85 global patterns, corresponding to 79 distinct TFs (Fisher Exact test, FDR<5%, *log*_10_(fold enrichment) > 1.5, Figure 2A).

To provide additional evidence that some of the TFs corresponding to these motifs potentially have trans-regulatory activity we analyzed their correlations with gene expression. In total, 24 of the 79 TFs with motifs enriched in a global pattern, also have their expression levels significantly associated with a pattern (FDR<5%, Figure 2C), although not necessarily the same pattern. Two TFs (TP73 and CREB5) had gene expression patterns associated with the same global pattern for which their motifs were enriched. TP73 has been shown to regulate T helper differentiation-related genes, which results in variation in autoimmune disease susceptibility in mice [44]. The CREB family of transcription factors regulate genes containing a cAMP-responsive element, including a number of immune-related genes [45]. TFs with expression associated with a global pattern, particularly the same one in which they have an enriched motif have additional evidence of trans-regulatory activity, but we note that TFs might be post-transcriptionally regulated, which could lead motif enrichments in specific global patterns without corresponding gene expression correlations.

### LCL global patterns increases power to detect trans-eQTLs

In traditional trans-eQTL testing, every variant is associated with the expression of every gene, resulting in a substantial multiple testing burden and low power to detect true associations. To mitigate this, an informed set of variants can be tested to reduce the multiple testing burden. For example, cis-eQTLs have been used as an informed set of variants because variants that affect expression locally are more likely to affect other genes indirectly [26].

We hypothesized that the gQTLs would be more likely to be trans-eQTLs than cis-eQTLs, since they are associated with global patterns of histone modifications that recur in many regions of the genome. Thus, we used gQTLs as an informed set of variants in trans-eQTL analysis using the LCL data set (n=75) and compared this with an approach using cis-eQTLs as an informed set of variants. We restricted the analysis to global patterns with signal in more than one individual to increase power of gQTL discovery (Methods), which led us to considering 62 gQTLs. We identified 45 significant trans-eQTLs (FDR < 5%) corresponding to 23 of 62 of the gQTLs when using the gQTLs as the informed set of variants. In comparison, an alternative approach of using cis-eQTLs as an informed set of variants did not have any significant trans-eQTLs (FDR < 5%).

We used data from the Whole Blood tissue of the GTEx data set (n=338) to attempt to replicate the LCL trans-eQTL identified. Of the 45 trans-eQTLs identified, only 10 could be tested for replication due to ancestry differences between the LCL data set and the GTEx data set (Methods). While none of these SNPs were individually significant after correcting for multiple testing (Bonferroni corrected *p* < 0.05), the p-values obtained in the replication analysis were more significant than we would expect by chance (Mann-Whitney U Test, *p* = 0.002, Methods). The lack of replication of individual SNPs after correcting for multiple testing is likely due to relatively small sample size of the replication cohort. Trans-eQTLs have been historically difficult to replicate due to small effect sizes and low power [46]. Our results suggest, that by using gQTLs instead of cis-eQTLs as an informed set of variants, we can be better powered for trans-eQTL discovery.

### Learning global patterns across autism spectrum disorder cases and controls

In order to directly associate global patterns with complex disease, we learned a separate model using H3K27ac histone modification data previously collected from prefrontal cortex tissue in ASD cases and controls [4].

We used the same procedures as the LCL data set to preprocess the data, train models, and select the number of hidden states except that we accounted for a larger set of known covariates that were used in previous differential peak analyses (Methods) [4]. We selected the 90-state model (Figure 4) for followup analyses, which maximized the number of gQTLs (Figure S9) with 2229 gQTLs. Similar to the LCL analysis, we performed GREAT [36] enrichment using the gQTLs as the foreground and the whole genome as the background. We identified a number of enriched GO terms (FDR < 5%, Supplementary Data 1) relevant for the prefrontal cortex tissue, such as “neuron projection morphogenesis”, “axon development”, and “regulation of axon guidance.” While we also observed significant enrichment for terms not directly related to brain function, we do not expect all global patterns to be cell type-specific.

**Figure 3:**
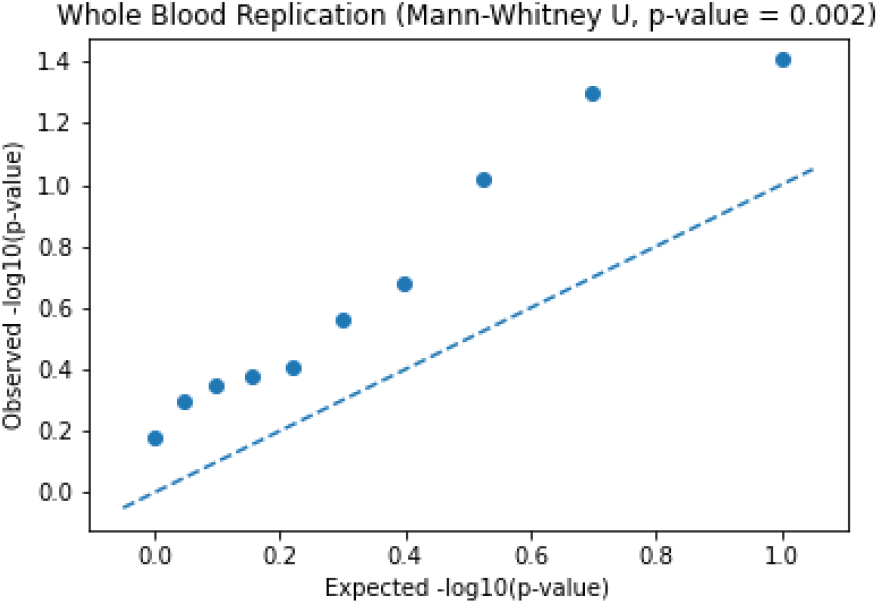
LCL trans-eQTL replication analysis. We performed a replication analysis on 10 trans-eQTLs identified in the LCL data with MAF of at least 5% in the GTEx data set. Expected −*log*10(*p*−*values*) (x-axis) were computed using theoretical values from a uniform distribution. Observed −*log*10(*p*−*values*) (y-axis) were computed in the replication analysis. The dashed line corresponds to the null, where p-values from the replication experiment have the same quantiles as those from a uniform distribution. The distribution of the replication p-values was significantly lower than we would expect by chance (Mann-Whitney U Test, *p* = 0.002).

**Figure 4:**
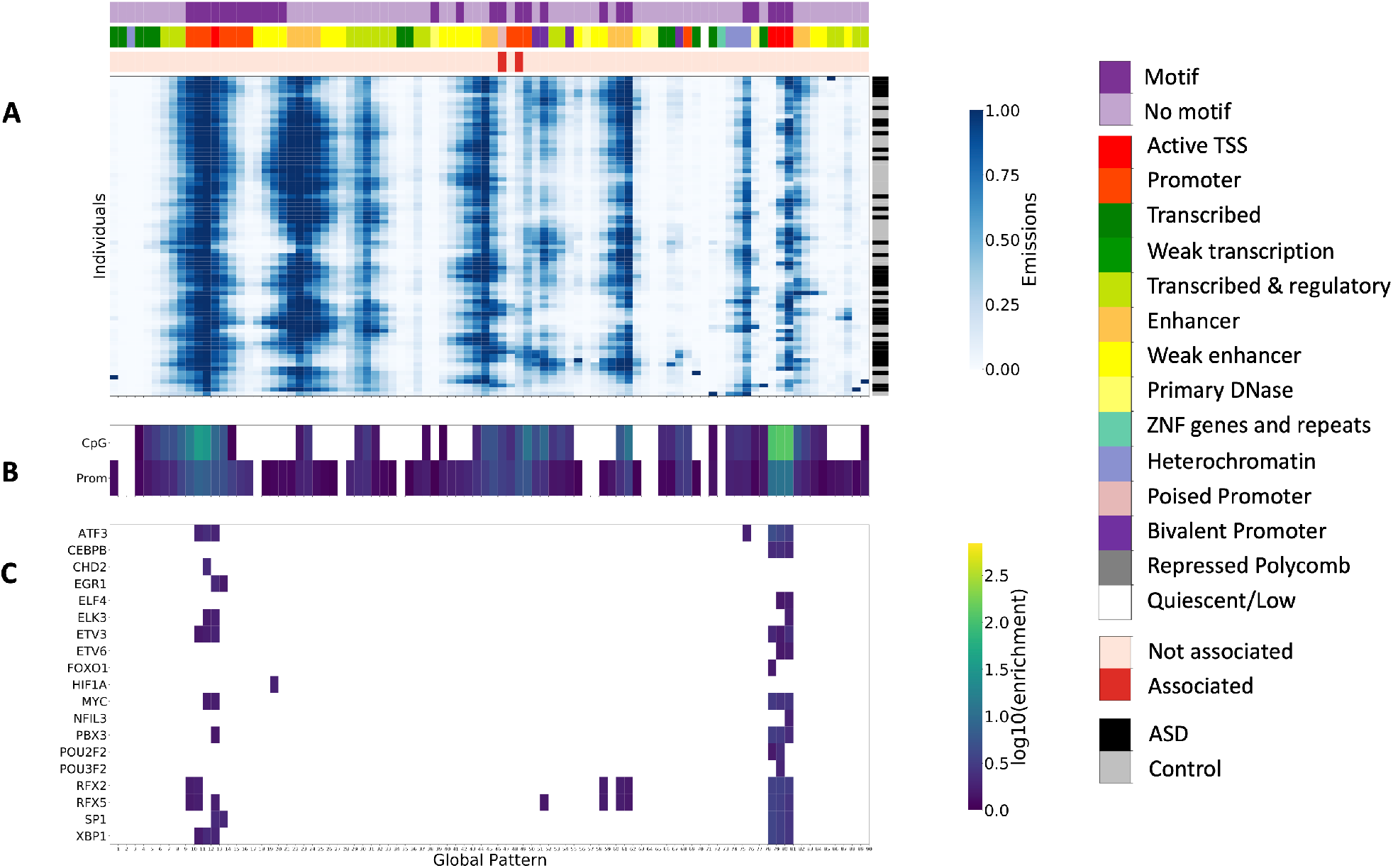
ASD 90-state model. A) Emission matrix for the 90-state ASD model. The x-axis (“Global Pattern”) shows the global patterns learned, and the y-axis shows each individual (“Individuals”). The diagnosis status of each individual is annotated (right). Each global pattern is annotated in the top three rows with whether the global pattern had significant motif enrichments (first), the reference chromatin state annotation for prefrontal cortex tissue [40] with the highest enrichment (second), and whether the state was significantly associated with diagnosis status (third). B) Overlap enrichments for CpG Islands [37] and promoter annotations computed from GENCODE TSS annotations [38] C) Motif enrichments for 19 TFs with motifs enriched in at least one global pattern (FDR<5%, log10(fold enrichment) > 1.5) and gene expression associated with global patterns (FDR<5%). Only significant enrichments are shown (Fisher’s Exact Test, FDR<5%, log10(enrichment) > 1.5).

**Figure 5:**
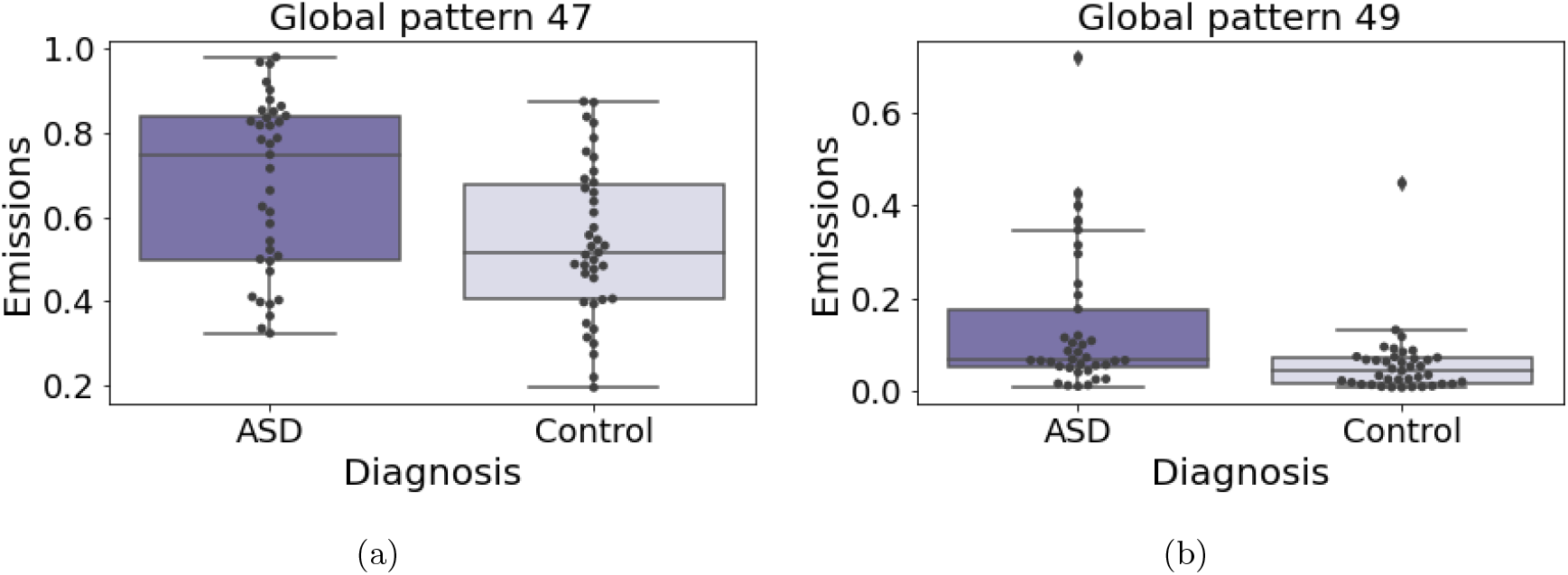
Association with ASD diagnosis status. We tested for association between emission parameters for each state in the 90-state ASD model and disease status. After controlling for multiple testing using a permutation test, two global patterns (Global pattern 47, left) and (Global pattern 49, right) were significantly associated with diagnosis status (*p* < .05, Mann-Whitney U Test). In each subplot, the diagnosis status is shown on the x-axis, and the emission parameters for the state are shown on the y-axis.

We next computed enrichments of global pattern for chromatin states previously annotated in the prefrontal cortex tissue from the Roadmap Epigenomics Consortium [47] (Figure S8). We annotated each global pattern with the most enriched state from this single reference epigenome annotation (Figure S8, Figure 4A). Like the LCL model, most of the states in the ASD model were highly enriched for promoter and enhancers (Binomial test, FDR<0.05) (Figure 4A). Consistent with the LCL data set, we also found high enrichments for CpG islands and GENCODE annotations of TSS in global patterns annotated as promoters (Figure 4B).

We identified 3535 genes with expression associated with the ASD global patterns (FDR<5%). In order to identify potential trans-regulators, we performed TF motif enrichment analysis for each global pattern. As with the LCL data, we calculated motif enrichment of 602 TF motifs in the ENCODE motifs database [43] compared to shuffled motifs. We observed enrichment of motifs for 27 of the 90 global patterns, corresponding to 114 distinct TFs (Fisher Exact test, FDR<5%, *log*_10_(fold enrichment)> 1.5, Figure 2A). In total, 19 of the 114 TFs with motifs enriched in a global pattern, also had their expression levels associated with the pattern (FDR<5%, Figure 2C), although not necessarily the same pattern. The motif enrichments for these TFs are shown in Figure 4C. For example, we found the RFX family’s motif, which was also previously identified to be enriched in differential peaks between cases and controls [4]. Furthermore, it was noted that RFX2 contains a differentially acetylated peak in its promoter, providing a potential mechanism by which its expression differs across individuals [4].

As the ASD data set contains both case and control individuals, we associated the global patterns with the diagnosis status. This identified two global patterns (47 and 49) that were significantly associated with ASD status (Mann-Whitney U, *p*_*adj*_ < 0.05, Methods). Both global patterns were highly enriched for promoter regions of the genome. Global pattern 47 was significantly associated with the expression of 22 genes, and global pattern 49 was significantly associated with the expression of 316 genes (FDR<5%). The genes associated with state 49 were enriched for several GO terms (Table S1), including “cellular response to interleukin-1,” which has been previously been suggested to play a role in the disease etiology [48]. These results suggest that analyzing global patterns can potentially be informative towards studying complex disease.

## Discussion

In this work, we learned global patterns of epigenetic variation across individuals and systematically annotated the human genome according to these patterns of variation using a stacked ChromHMM model. We applied this framework to an LCL data set for three histone modifications (H3K27ac, H3K4me1, H3K4me3) and an ASD case and control data set for one histone modification (H3K27ac).

Previous work in detecting molecular variation across individuals have performed marginal association tests on consensus regions of the genome [1, 2, 3, 4, 5]. However, these analyses do not investigate whether the patterns of variation across individuals recur in multiple regions of the genome. By associating genes and variants with the global patterns, we decrease the multiple testing burden substantially for detecting trans-regulators and trans-eQTLs. Thus, this investigation is better powered than traditional association methods that consider larger numbers of potential associations.

We identified motif of TFs enriched in the global patterns for both the LCL and ASD data enabling us to predict potential trans-regulators. Of these TFs, a fraction also had differential expression associated with at least one global pattern. In these cases, there is additional evidence of molecular coordination in different regions of the genome. We also identified thousands of genes in both data sets with expression patterns significantly associated with the global patterns.

We also utilized the global patterns to improve power of trans-eQTL studies. By using variants associated with the global patterns as an informed set of variants for testing, we identified 45 trans-eQTLs in the LCL data set (FDR<5%). We performed a replication analysis for 10 of these variants with MAF > 5% in the GTEx data set, and observed that the replication was significantly higher than expected by chance. Since the sample sizes of both the LCL data set (n=75) and the GTEx Whole Blood Tissue (n=338) limited the statistical power in both the initial and replication study.

One challenge in identifying global patterns is that it is difficult to distinguish global patterns due to confounders and those due to biological reasons, such as trans-regulation. Unsupervised methods, such as principal component analysis and PEER [49] are likely to remove true biological signal in these applications. To mitigate the effect of confounders, we regressed out known covariates from the histone modification input signal. In the LCL data set, we showed that the biological signal of interest was consistent across multiple histone modifications. To further demonstrate the likely biological significance of the global patterns we relied on external data sets and annotations. For both data sets, we identified a number of global patterns that showed consistent co-variation with gene expression. We also showed that the global patterns were associated with common genetic variants and that these variants were in close proximity to tissue/condition relevant genes. These analyses support the differences across individuals captured across individuals is biological. However, we cannot exclude that technical differences are also driven by sample-level confounding factors that were not previously regressed out.

Finally, we identified two global patterns significantly associated with ASD, in spite of the small sample size and heterogenous nature of ASD. The framework we have used is general and can be applied to other datasets to further analyze and understand epigenetic variation across individuals and its relationship with complex disease.

## Methods

### Histone modification data

We learned global patterns in LCL histone modification data [15] and ASD histone modification data [4]. In the LCL data, histone modification signal was mapped for three marks (H3K27ac, H3K4me1, H3K4me3) in 75 individuals. In the ASD data, H3K27ac signal was mapped in the prefrontal cortex tissue for 93 individuals. We used the same 76 individuals that were used previously after quality control [4]. Only autosomes were included in this analysis and we used the hg19 genome assembly.

### Learning epigenetic patterns across individuals using a stacked ChromHMM model

We quantified the number of reads falling within each 200bp non-overlapping bin in the genome using the BinarizeBam command in the ChromHMM software package (version 1.11) [21]. For each bin, we then fit a quasi-poisson regression model, where the dependent variable was the number of reads in the bin and the independent variables were standardized known covariates [50]. Quasi-poisson regression handles both over and under-dispersion of count data and has been used in related applications [51]. We quantile normalized the read counts so that the distribution of counts across bins in the genome was the same for each individual. We regressed out the effect of the known covariates for each data set that were used in previous work. For the LCL data set, we accounted for sex, genotyping method, number of reads and relative strand correlation (RSC) [15]. We performed the correction separately for each mark in the LCL dataset. For the ASD data set, we accounted for age, sex, percentage of neuronal cells, brain bank, number of peaks, fraction of reads in peak (FRIP), read duplicate fraction, and read alignment fraction [4]. We then binarized the corrected histone modification count data according to the procedures used in ChromHMM using the BinarizeSignal command with default parameters [21]. We used binarized read count data as input to learn the parameters to a stacked ChromHMM model. To learn the model parameters and perform genome segmentation and annotation we used the LearnModel command of the ChromHMM software package with default parameters [21]. We trained “stacked” models where each ChIP-seq dataset is treated as a separate mark. We trained separate models using 5-100 hidden states in increments of 5 states for each data set.

### gQTL analysis

We associated each global pattern with common variants (MAF > 5%) previously identified in each data set [15, 4]. In this association analysis, we split the emission matrix by mark and treated the emission parameters for each mark and state as a phenotype. We performed the associations and computed the p-values for each association using the MatrixEqtl software [52]. We use a Bonferonni corrected threshold of 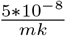, where *m* in the number of marks, and *k* is the number of global patterns. For the LCL data set, we performed the gQTL analysis on all 75 individuals with Yoruban ancestry. For the ASD data set, we performed the gQTL analysis using on individuals with Caucasian ancestry (69/76 individuals).

### Choosing number of hidden states

To select the number of hidden states, we first trained models using between 5 and 100 hidden states in increments of 5 states. We then associated common variants (MAF>5%) in the human genome with the emission parameters to identify gQTLs and chose the number of hidden states that maximized the number of gQTLs.

### Evaluating robustness of global patterns

To evaluate whether the global patterns are robustly learned across different subsets of data, for each number of states between 5 and 100, we first trained two models using chr1 and chr2 separately. For each pair of models with the same number of states, we matched states from the two models using a greedy approach. We iteratively computed pairwise Spearman correlations between unpaired states, matching the two states from the models with the highest correlation.

### GREAT enrichment analysis

We performed GREAT enrichment [36] for the set of gQTLs identified by each data set, using the gQTLs as the foreground and the whole genome as the background. The whole genome was selected as the background distribution because variants from all regions of the genome were tested for gQTL association. Using default parameters, GREAT computes a basal regulatory domain of a minimum distance of 5kb upstream and 1kb downstream. The regulatory domain is extended in both directions to the nearest gene’s basal domain, but no more than 1000kb in one direction. gQTLs are intersected with these regulatory domains to identify genes in close proximity to the gQTLs. GREAT uses a hypergeometric test to assess whether the foreground is significantly enriched for Gene Ontology (GO) terms relative to the background. We identified GO terms that were significantly enriched in the gQTLs (FDR<5% and observed genes > 5).

### Overlap enrichments

We performed overlap enrichments for external annotations in both the LCL and ASD models using the OverlapEnrichment command in the ChromHMM software (Version 1.11) [21]. To assess the significance of the overlap, we computed significance p-values for the enrichments using a binominal test, where the probability of success was set the the fraction of bases covered by the annotation in the genome. We corrected for multiple testing using an FDR threshold of 5% (Benjamini-Hochberg procedure) for all annotations tested within each model.

For both models, we computed enrichments for tissue-specific chromatin state annotations using a previous 25-state “concatenated model” that was learned from imputed data for 12-marks [40] based on data from the ENCODE Consortium Project [53] and the Roadmap Epigenomics Project [47]. For the LCL model, we used the lymphoblastoid cell line (reference epigenome E116), and for the ASD model, we used the prefrontal cortex tissue (reference epigenome E073). We tested enrichment for each of the 25 states in these models and annotated the global patterns with the state with the highest enrichment. We also computed enrichments for promoter annotations computed from GENCODE V29 TSS annotations [38], which were defined as regions 2KB from the TSS of the genes. Finally, we computed enrichments for CpG Islands obtained from the UCSC Genome Browser [54].

For the LCL model, we computed enrichments for DNase I hypersensitive sites collected in the same individuals [15]. Specifically, we tested previously published consensus peaks that were obtained by merging peaks across individuals (http://mitra.stanford.edu/kundaje/portal/chromovar3d) [15]. We also obtained 4,950 candidate causal SNPs for 21 immune diseases that were previously fine-mapped from 636 autoimmune GWAS loci [9] and tested them for enrichment in the global patterns.

### Association of global patterns with gene expression and protein quantification

For the association of global patterns with gene expression, we used external RNA-seq data that was not used to train the model. Expression data was previously collected for 54 of the 75 LCL samples using RNA-seq [15]. We corrected the expression data for known covariates (sex, genotyping method, number of reads, the ratio between the fragment-length peak and the read-length peak or relative strand correlation (RSC) [55]). We only considered genes with variability as defined in a previous publication [15]. Expression data was available for 51 of the 76 individuals from the ASD data set [30]. Expression values were corrected for known covariates as described in [30]. We only considered genes with variability as defined in this previous work. We tested for association between gene expression and the emission parameters. We calculated the association statistics for a linear model using the MatrixEqtl software [52] and used an FDR threshold of 0.05 within each mark.

For the LCL data set, we obtained log2 protein quantification data for 60 of the 75 individuals for a subset of 4371 genes with both gene expression and protein quantification data [41]. Using the same association framework as the association between global patterns and gene expression, we associated the log2 protein quantification with the global patterns for each mark separately. We used an FDR threshold of 0.05 within each mark.

### Motif enrichments

We computed enrichment of 602 TF motifs in the ENCODE motifs database using a tool published with the database [43]. Briefly, this tool computes the enrichment of each motif compared to a background created using shuffled control motifs with a confidence interval correction for small counts. Although this database includes both known and computationally discovered motifs, only known motifs, which are more specific to individual TFs, were used. We identified motifs with FDR<5% and a log2 fold enrichment of at least 1.5.

### Trans-eQTL analysis

We performed two trans-eQTL analyses using the LCL dataset. First, we performed a baseline trans-eQTL analysis, where cis-eQTLs identified in prior studies [15] were tested for association with all expressed genes, as defined previously for associations with gene expression and emission parameters. We compared this trans-eQTL analysis to one where gQTLs were tested for association with all expressed genes. For both analyses, we used an FDR threshold of 5%.

For the trans-eQTL analysis using gQTLs as the informed set of SNPs, we further restricted the set of SNPs to those that were associated with “nonsingleton global patterns” to increase power. This is analogous to filtering out SNPs with low minor allele frequency and would theoretically improve power for downstream replication. We identified nonsingleton global patterns as follows. For each global pattern, we counted the number of individuals with emission parameters greater than 0.5 within that global pattern. We defined singleton states as those with only one individual with emission parameter greater than 0.5. All other states were classified as nonsingleton states, including states where all individuals had emissions less than 0.5.

For the LCL trans-eQTLs identified using gQTLs as the informed set of SNPs, we performed a replication analysis using the Whole Blood Tissue from the GTEx Consortium v6p [56]. Only 10 SNPs could be tested for replication due to differences in ancestry between the LCL data set (Yoruban) and the GTEx data set (primarily European). In this analysis, we used the same covariates as used for eQTL discovery in GTEx [26] (3 genotype principal components, 135 PEER factors, genotyping platform, and sex). To compute the replication between each significant gQTL and gene pair, we used a Bonferroni corrected threshold of 0.05. We compared the p-values obtained in the GTEx replication with a uniform distribution using a Mann Whitney U statistical test.

### Association of global patterns with ASD disease status

We associated each global pattern in the 90-state ASD model with the disease status using a Mann-Whitney U test. We performed a permutation test to correct for multiple testing, where for each permutation, the samples labels were shuffled and the minimum p-value was retained for the null distribution. The adjusted p-values were computed as the fraction of p-values from the null distribution that were more significant than the observed p-values. We used an adjusted p-value threshold of 0.05 to identify states significantly associated with disease status.

## Data availability

Our work integrates data from a number of previous publications, including LCL ChIP-seq and RNA-seq data (http://mitra.stanford.edu/kundaje/portal/chromovar3d/) [15], LCL protein data [41], ASD ChIP-seq data (https://www.synapse.org/#!Synapse:syn4587616) [4], and ASD RNA-seq data (https://www.synapse.org/#!Synapse:syn11242290) [30].

The GREAT enrichments for gQTLs are available in Supplementary Data 1. The model parameters and segmentations learned in this paper are available in Supplementary Data 2 and Supplementary Data 3 for LCL and ASD, respectively.

## Acknowledgements

We would like to acknowledge Daniel Geschwind and Gokul Ramaswami for helping us access the ASD data and for providing feedback on the results.

This work was supported in part by the United States National Institutes of Health (DP1DA044371, R01MH109912, U01MH105578, UH3NS104095, U01HG012079) (J.E.), a Rose Hills Innovator Award (J.E.), the UCLA Jonsson Comprehensive Cancer Center and Eli and Edythe Broad Center of Regenerative Medicine and Stem Cell Research Ablon Scholars Program (J.E.), J.Z. received supported by a National Science Foundation Graduate Research Fellowship under Grant DGE-1650604 (J.Z.) and National Institutes of Health Award Number T32MH073526 (J.Z.).

## Supplementary information

**Figure S1:**
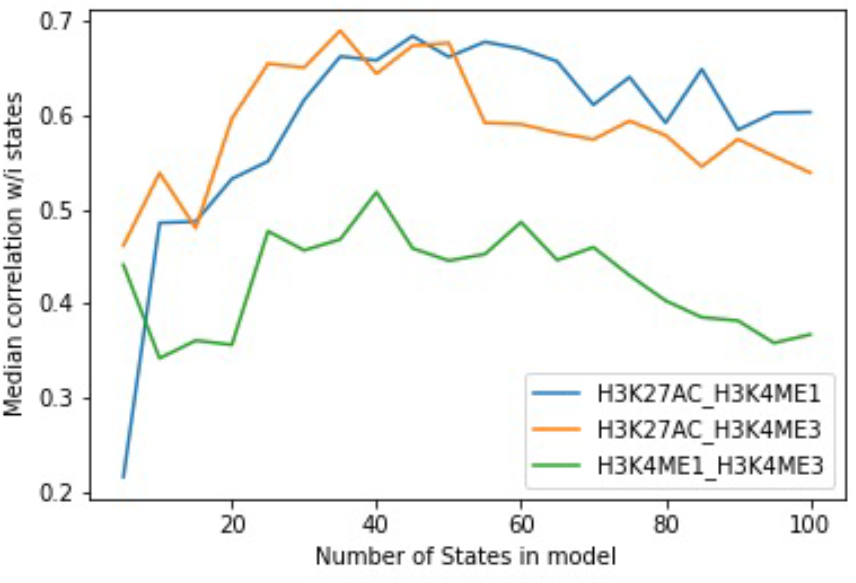
Pairwise correlation of marks in LCL data set. For each pair of histone modifications (H3K27ac, H3K4me1, H3K4me3), we computed the median Spearman correlation of the emission parameters across individuals within each state. Each colored line corresponds to a pair of marks. The median pairwise correlation (y-axis) is shown as a function of the number of states used to train each model (x-axis).

**Figure S2:**
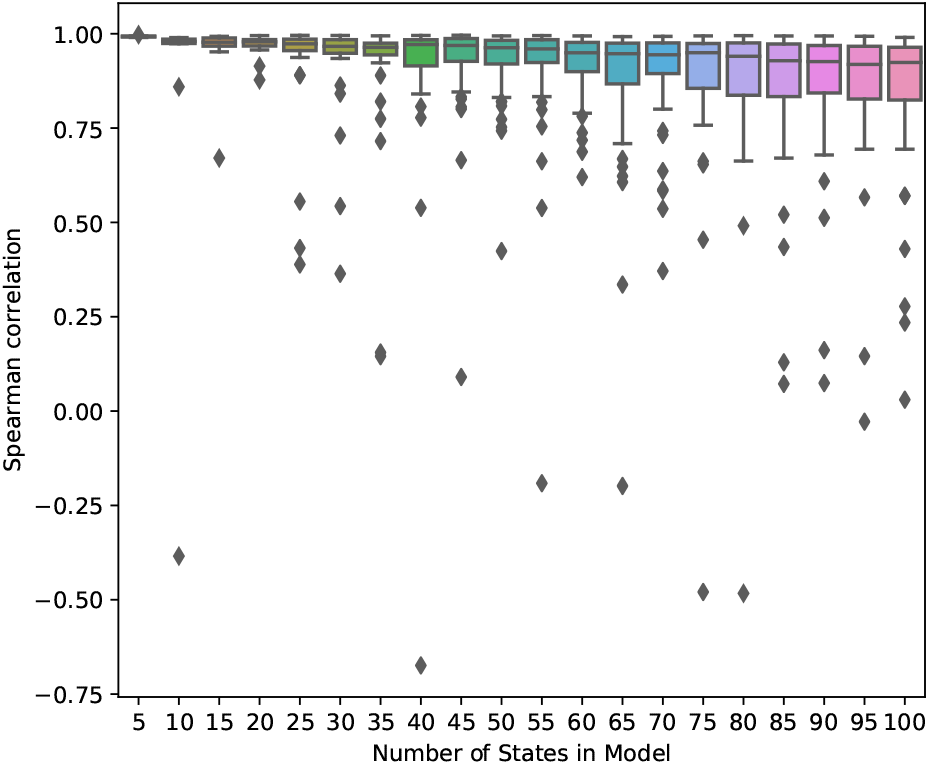
Pairwise state correlations in LCL data set. For each number of states (x-axis), two models were trained on different subsets of data (Methods). Each dot corresponds to the Spearman correlation (y-axis) of the emission parameters of a state in one model with a paired state in the other model. States between the two models were paired using a greedy algorithm (Methods).

**Figure S3:**
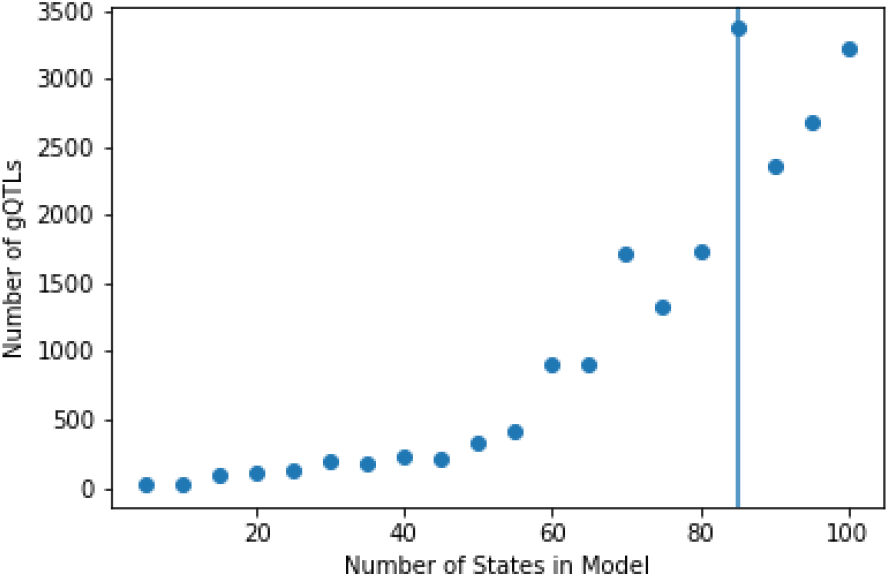
LCL model gQTLs. The total number of significant gQTLs (FDR<5%) in the LCL model (y-axis) is shown as a function of the number of states (x-axis). The number of gQTLs is maximized using an 85-state model denoted with the vertical line.

**Figure S4:**
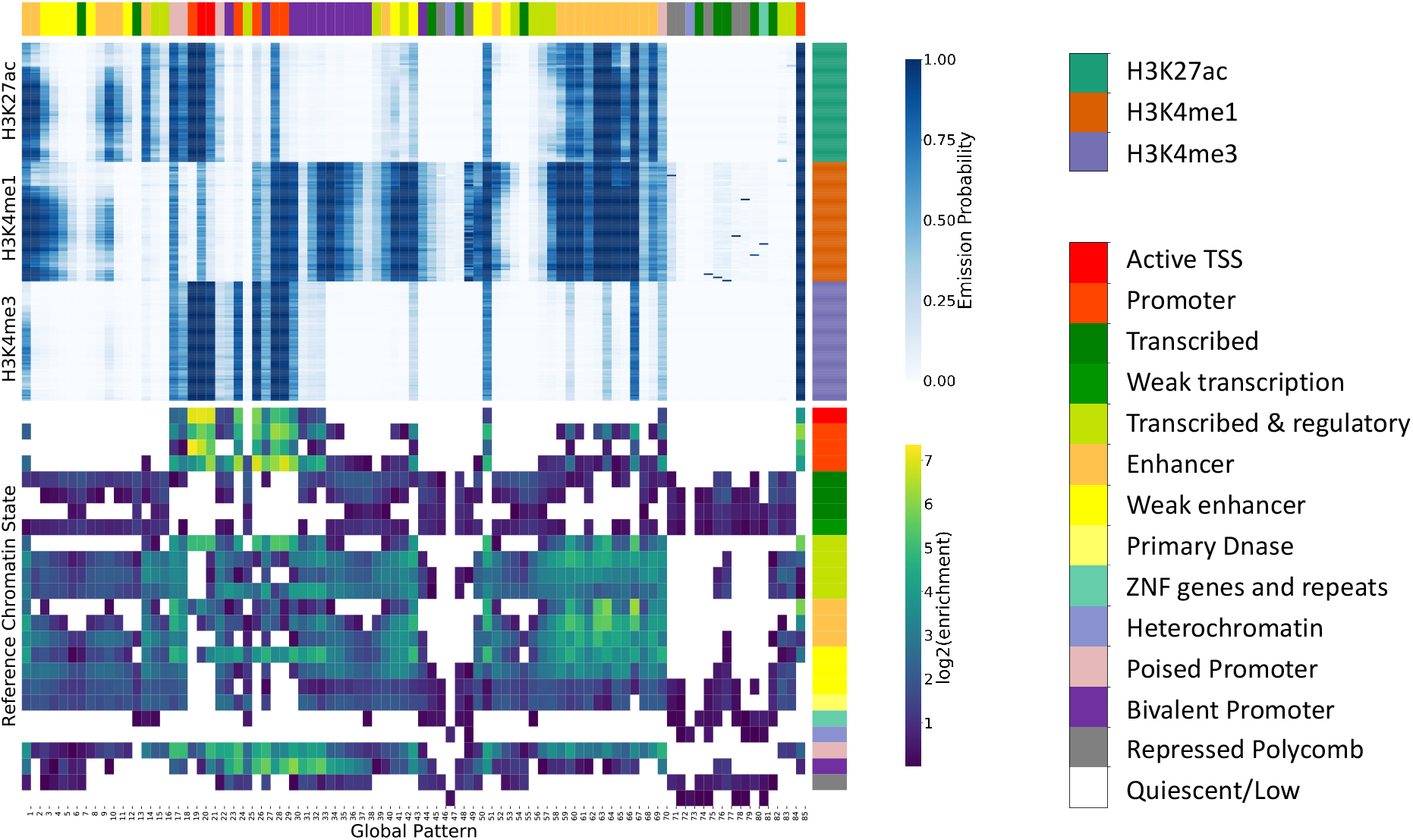
Fold enrichment of chromatin states for LCL 85-state model. The top heatmap shows the emission parameters learned for the 85-state model using the LCL dataset. The global patterns are on the x-axis, and the datasets are on the y-axis. The datasets are grouped by histone modification, and the individuals have the same ordering within each mark. The bottom heatmap shows the log2 fold enrichment of a previous reference chromatin state annotation in LCL for one individual based on imputed data for 12-marks, where states with the same annotation color (left) represent different sub-states of the same category [40]. Significant enrichments are indicated with color (Binomial Test, FDR<5%). The global patterns are shown on the x-axis (“Global Pattern”), and the LCL reference chromatin states on the y-axis (“Reference Chromatin State”). Each global pattern is annotated with the highest enriched chromatin state from the reference individual (top).

**Figure S5:**
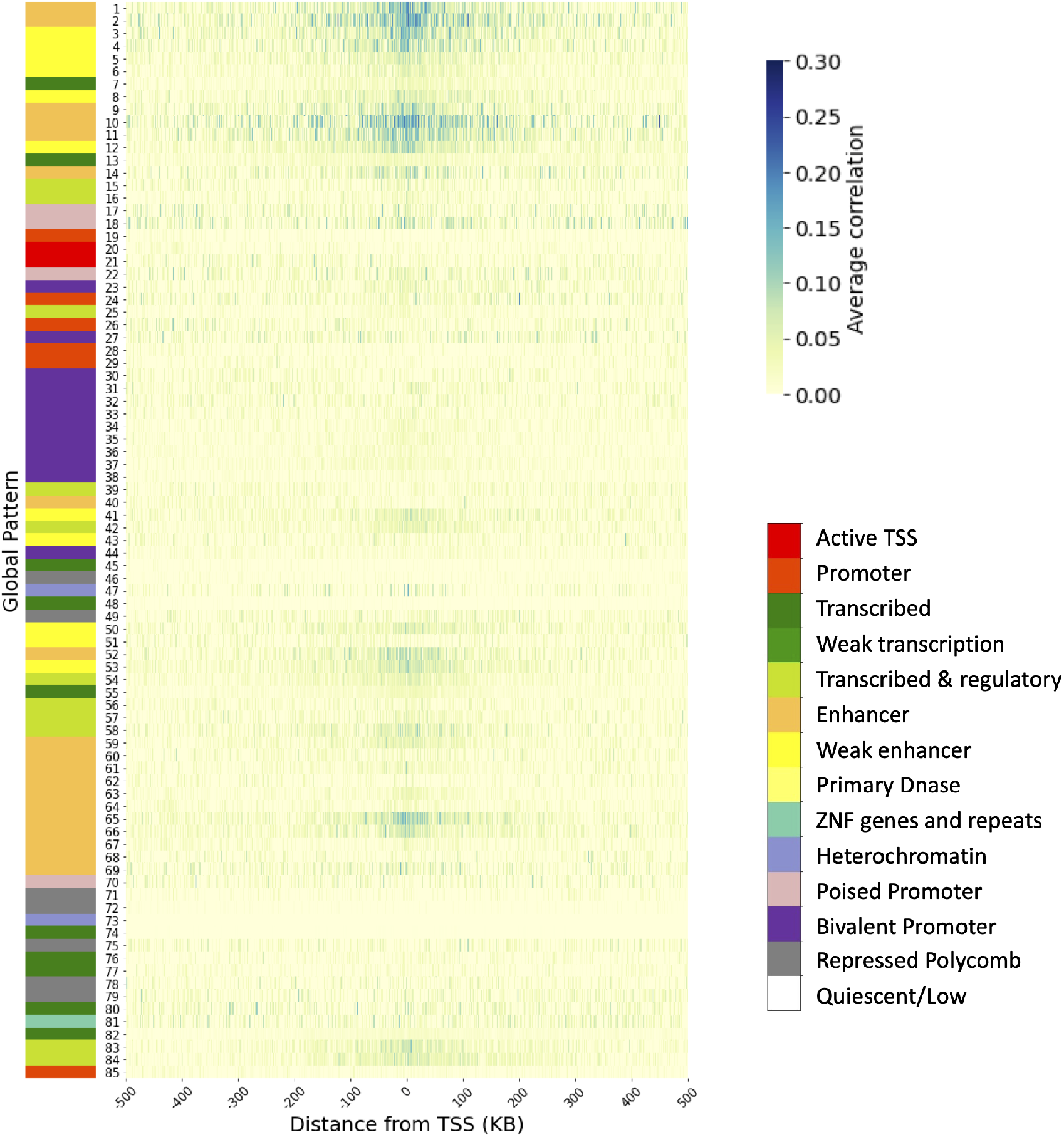
Average correlation of H3K27ac LCL emission parameters and gene expression as a function of distance. The genome segmentation based on the LCL global patterns was used to identify genes with transcription start sites (TSS) within 500KB for each global pattern. These genes were associated with the global pattern. The average Pearson correlations between emission parameters and expression of genes are shown for each global pattern (y-axis) using only genes with TSS within each 100bp bin (x-axis). The LCL reference chromatin state [40] with the highest enrichment for each global pattern is indicated based on the color on its left and the color legend on right.

**Figure S6:**
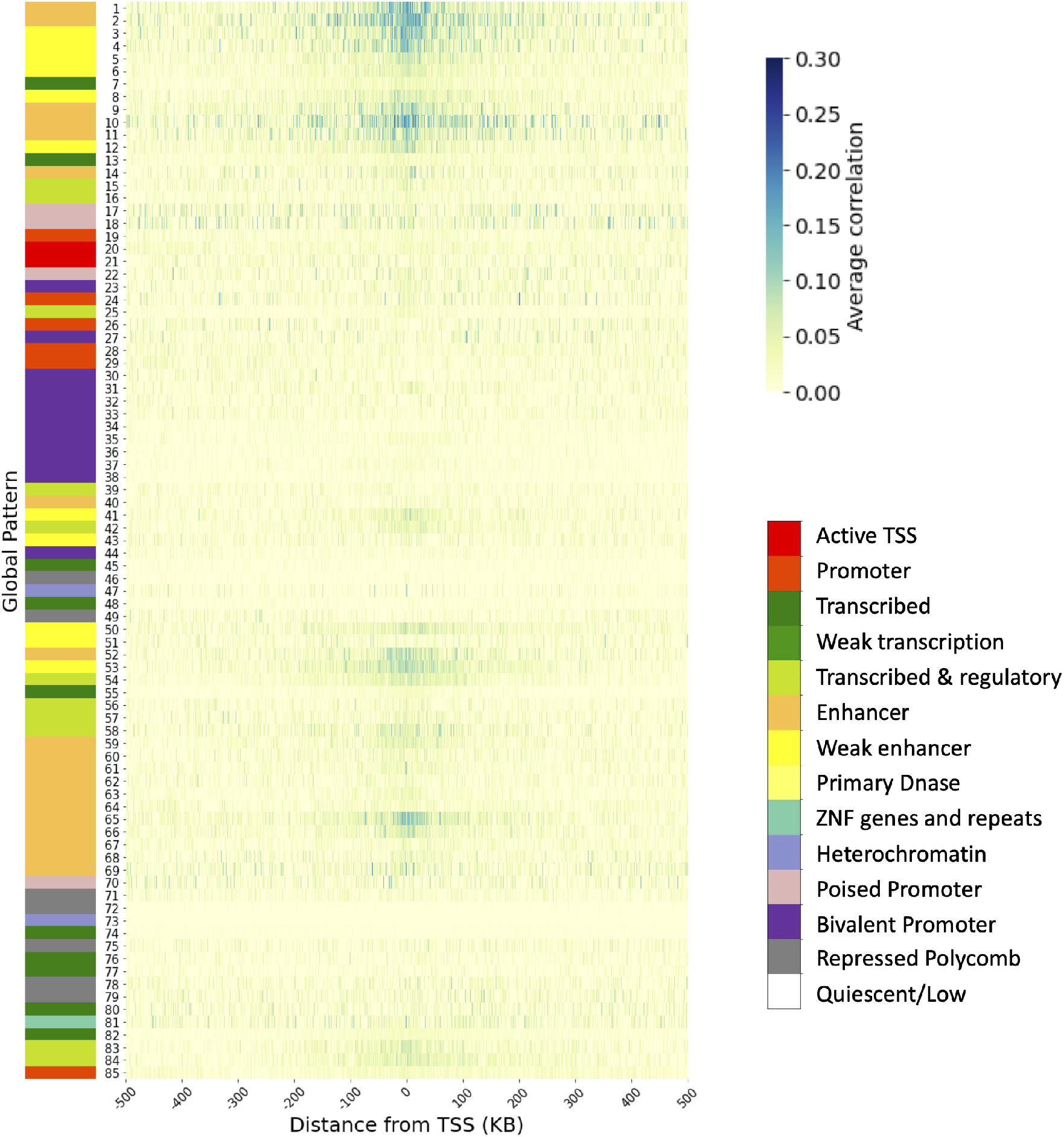
Average correlation of H3K4me1 LCL emission parameters and gene expression as a function of distance. Average correlation of H3K4me1 LCL emission parameters and gene expression as a function of distance. The genome segmentation based on the LCL global patterns was used to identify genes with transcription start sites (TSS) within 500KB for each global pattern. These genes were associated with the global pattern. The average Pearson correlations between emission parameters and expression of genes are shown for each global pattern (y-axis) using only genes with TSS within each 100bp bin (x-axis). The LCL reference chromatin state [40] with the highest enrichment for each global pattern is indicated based on the color on its left and the color legend on right.

**Figure S7:**
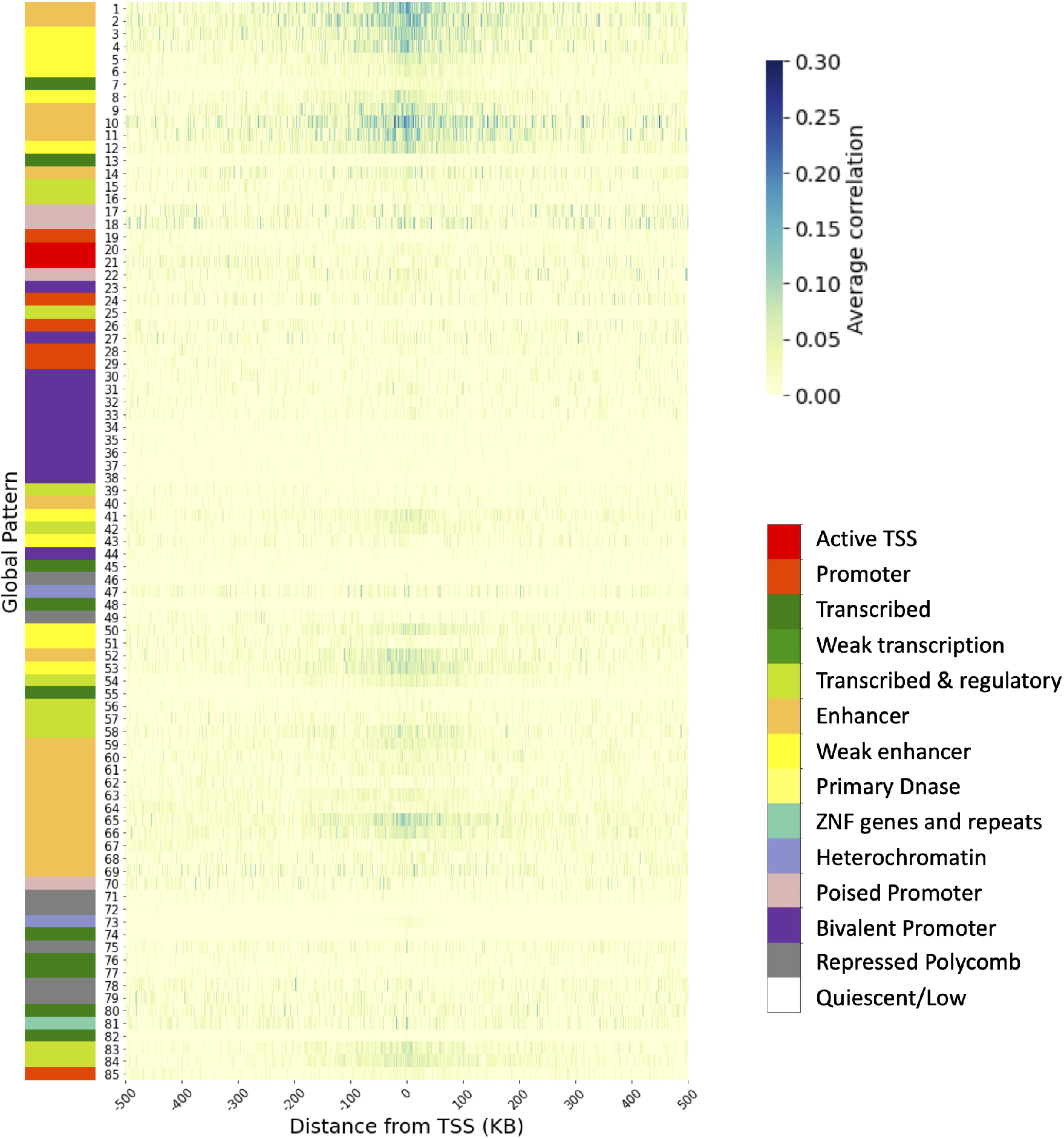
Average correlation of H3K4me3 LCL emission parameters and gene expression as a function of distance. Average correlation of H3K4me3 LCL emission parameters and gene expression as a function of distance. The genome segmentation based on the LCL global patterns was used to identify genes with transcription start sites (TSS) within 500KB for each global pattern. These genes were associated with the global pattern. The average Pearson correlations between emission parameters and expression of genes are shown for each global pattern (y-axis) using only genes with TSS within each 100bp bin (x-axis). The LCL reference chromatin state [40] with the highest enrichment for each global pattern is indicated based on the color on its left and the color legend on right.

**Figure S8:**
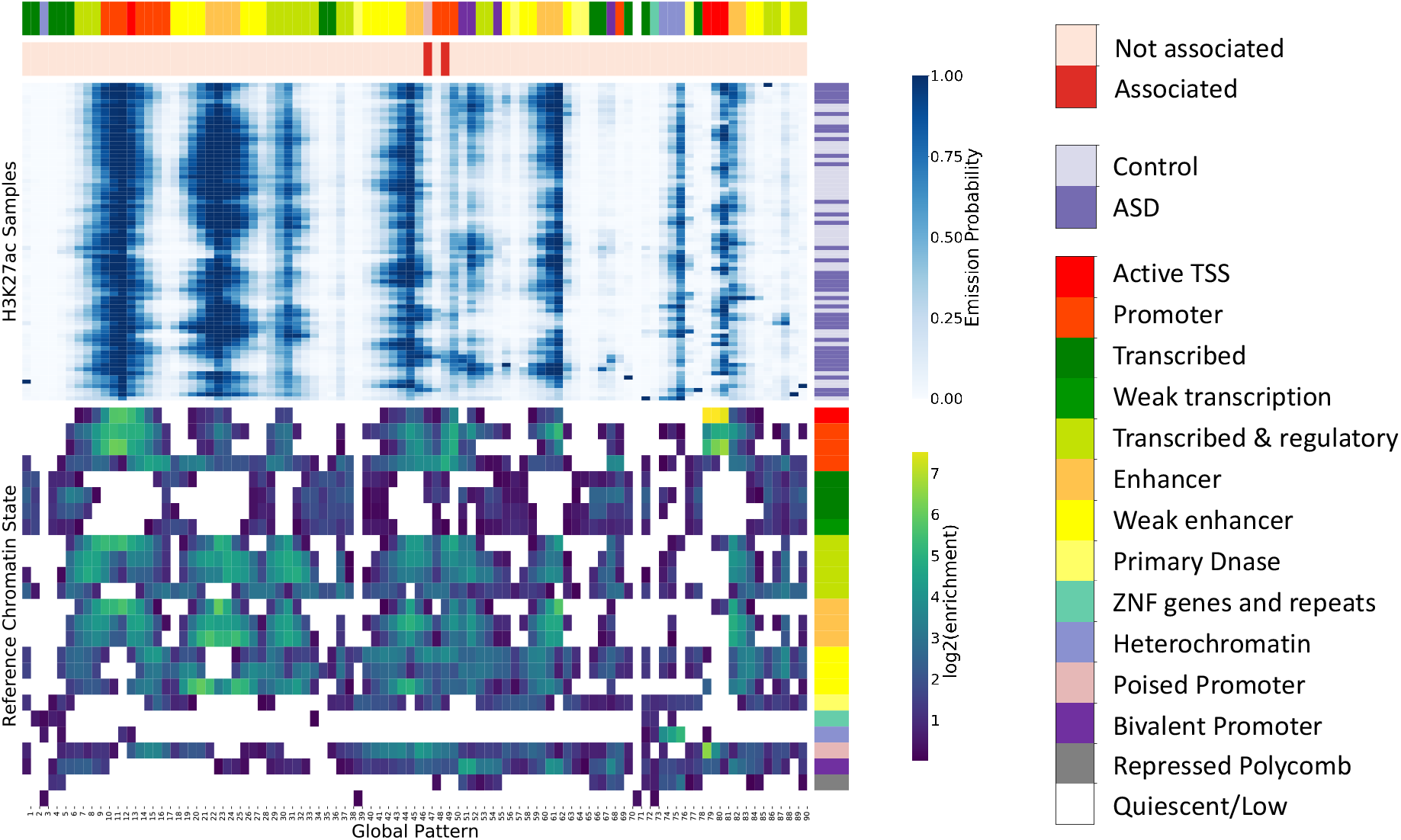
Fold enrichment of chromatin states for ASD 90-state model. The top heatmap shows the emission parameters learned for the 90-state model using the ASD dataset. The global patterns are on the x-axis. The samples are on the y-axis and are annotated as being either a case (dark purple) or a control (light purple) to the right. The bottom heatmap shows the log2 fold enrichment of a previous chromatin state annotation of a prefrontal cortex for a single reference epigenome using imputed data for 12-marks [40] (“Reference Chromatin State”). Significant enrichments are indicated with color (Binomial Test, FDR<5%). The global patterns are annotated with two tracks above the emission parameters. The first shows the chromatin state from the reference prefrontal cortex chromatin state annotation with the highest enrichment. The second shows whether the state is significantly associated with ASD status (red) or not (pink).

**Figure S9:**
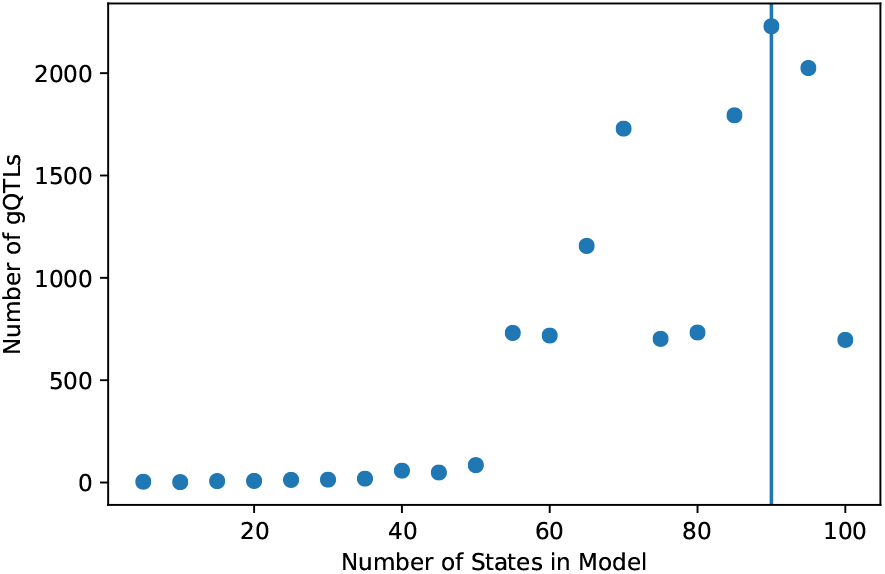
ASD model gQTLs. The total number of significant gQTLs (FDR<5%) is shown (y-axis) as a function of the number of states in the model (x-axis). Each point corresponds to one stacked ChromHMM model. The number of gQTLs is maximized using an 90-state model denoted with the vertical line.

**Table S1:** GO enrichment of genes associated with global pattern 49. We performed GO enrichment of the 316 genes associated with global pattern 49 using the whole genome as the background. The terms shown (“GO term”) were significantly enriched after correcting for multiple testing using an FDR threshold of 5%. The total number of genes for the term (“Total”), the observed number of genes in the foreground (“Obs”), the expected number of genes (“Exp”), the fold enrichment (“Fold Enrichment”), the raw p-value (“P”), and the FDR corrected p-value (“FDR”) are shown in the table above.

| **GO term** | **Total** | **Obs** | **Exp** | **Fold Enrichment** | **P** | **FDR** |
| --- | --- | --- | --- | --- | --- | --- |
| cellular response to interleukin-1 | 114 | 9 | 1.53 | 5.89 | 3.85E-05 | 3.01E-02 |
| rRNA processing | 219 | 12 | 2.94 | 4.09 | 6.27E-05 | 4.46E-02 |
| macromolecule modification | 2922 | 65 | 39.17 | 1.66 | 4.04E-05 | 3.02E-02 |
| protein metabolic process | 3900 | 82 | 52.28 | 1.57 | 1.92E-05 | 1.68E-02 |
| cellular macromolecule metabolic process | 4497 | 92 | 60.28 | 1.53 | 1.37E-05 | 1.27E-02 |

